# The expression of YAP1 is increased in high-grade prostatic adenocarcinoma but is reduced in neuroendocrine prostate cancer

**DOI:** 10.1101/832360

**Authors:** Siyuan Cheng, Nestor Prieto-Dominguez, Shu Yang, Zachary M. Connelly, Samantha StPierre, Bryce Rushing, Andy Watkins, Lawrence Shi, Meredith Lakey, Lyndsey Buckner Baiamonte, Tajammul Fazili, Aubrey Lurie, Eva Corey, Runhua Shi, Yunshin Yeh, Xiuping Yu

## Abstract

**BACKGROUND:** After long-term androgen deprivation therapy, 25-30% prostate cancer (PCa) acquires an aggressive neuroendocrine (NE) phenotype. Dysregulation of YAP1, a key transcription coactivator of the Hippo pathway, has been related to cancer progression. However, its role in neuroendocrine prostate cancer (NEPC) has not been assessed.

**METHODS:** Immunohistochemistry was used to evaluate YAP1 protein levels during PCa initiation and progression. YAP1 knockdown and luciferase reporter assays were used to evaluate the ability of YAP1 to modulate Wnt/beta-Catenin signaling.

**RESULTS:** YAP1 expression was present in the basal epithelial cells in benign prostatic tissues, lost in low grade PCa, but elevated in high grade prostate adenocarcinomas. Interestingly, the expression of YAP1 was reduced/lost in both human and mouse NEPC. Finally, YAP1 knockdown in PCa cells activates Wnt/beta-Catenin signaling, which has been implicated in NE differentiation of PCa, supporting a functional involvement of the loss of YAP1 expression in NEPC development.

**CONCLUSIONS:** The expression of YAP1 is elevated in high grade prostate adenocarcinomas while lost in NEPC. Reduced YAP1 activates Wnt/beta-Catenin signaling in PCa cells. These results suggest that when applied to PCa patients, YAP1 inhibitors shall be used with caution.

## INTRODUCTION

Prostate cancer (PCa) is the most commonly diagnosed cancer among American men.^1^ Although androgen deprivation is an effective therapy for advanced and recurrent PCa, most of the prostate tumors eventually become resistant to treatment and progress to castrate-resistant prostate cancer (CRPC).^2^ In the disease progression, PCa cells may lose the prostatic adenomatous features and trans-differentiate into neuroendocrine prostate cancer (NEPC).^3^ NEPC is an aggressive pathological type that lacks the responsiveness to androgen deprivation tharepy. The survival of the patients with NEPC has been markedly reduced in comparison to adenocarcinomas. Therefore, there is a pressing need for studying the molecular mechanisms that drive the NE differentiation of PCa cells and developing novel agents that could effectively suppress the emergence of this lethal phenotype.

Wnt/β-Catenin signaling is involved in PCa progression. Activation of this pathway promotes the development of high-grade prostate intraepithelial neoplasia (PIN) in transgenic mice^4^. Combined activation of Wnt/β-Catenin signaling and SV40 T-antigen results in the development of invasive prostatic adenocarcinoma (AdPCa) with increased NE differentiation.^5^ Moreover, activation of Wnt/β-Catenin induces the expression of FOXA2, a transcription factor that is expressed in both human and mouse NEPC.^5^ These results indicate that Wnt/β-Catenin signaling is involved in NEPC tumorigenesis. However, it is not clear how Wnt/β-Catenin signaling is activated during PCa progression. In this study, we showed that the expression of YES-associated Protein (YAP1) was increased in high-grade AdPCa but was lost in NEPC, and decreased YAP1 expression activated Wnt/β-Catenin signaling, implicating YAP1 in the activation of this signaling pathway and the acquisition of NE phenotype during PCa progression.

YAP1 and its homologous protein TAZ are key transcriptional coactivators involved in the modulation of several processes in mammalian cells, such as glucose uptake, proliferation, spreading, apoptosis and differentiation.^6^ The activity of these transcriptional coactivators is mainly modulated by the Hippo pathway,^6^ which when activated impairs the translocation of YAP1 and TAZ to the nucleus and promotes their proteasome-mediated degradation.^6^ Alternatively, when the Hippo pathway is inactivated, YAP1/TAZ can enter the nucleus and activate the transcription of target genes.^6^ Additionally, their translocation to the nucleus can be modulated by Wnt/β-Catenin signaling in a Hippo-independent manner,^7^ and the loss of YAP1/TAZ can induce β-Catenin-dependent gene expression, ^7^ suggesting that the Hippo and the Wnt/β-Catenin pathways are interconnected to develop an integrated response.^8^

Dysregulation of YAP1 has been related to cancer initiation and progression.^9,10^ In CRPC, YAP1 are associated with cancer proliferation and invasiveness.^11^ However, the significance of YAP1 in NEPC has not been evaluated. In the present study, we found that the expression of YAP1 was down regulated in NEPC, and we also showed that downregulation of YAP1 activated Wnt/β-Catenin signaling, which has been previously shown to promote NE differentiation of PCa,^5,12^ supporting a role of loss of YAP1 in NEPC development. To the best of our knowledge, this is the first study demonstrating a biological role of YAP1 in NEPC development.

## MATERIAL AND METHODS

### Sample collection

De-identified human prostate tissue specimens were used in accordance with LSU Health-Shreveport IRB protocols. TRAMP, 12T-10 LADY, and NE10 archived tumor sections were used for this study.

### Immunohistochemistry and Immunofluorescence staining

Immunostaining was performed using Vectastain elite ABC peroxidase kit (Vector Laboratories, Burlingame, CA) as described previously.^13^ Primary antibodies of YAP1 and Chromogranin A (CHGA) were purchased from Santa Cruz Biotechnology (Dallas, TX), Synaptophysin (SYP) (BD biosciences, San Jose, CA), p63 and FOXA2 (Abcam, Cambridge, MA). The tissue sections were counterstained, mounted, and imaged with a Zeiss microscope (White Plains, NY). The percentage of cells stained was evaluated on a scale of 1+ (1-25%), 2+ (25-50%), 3+ (50-75%), and 4+ (75-100%) and the intensity of expression, 0 (negative), 1+ (weak), 2+ (moderate), and 3+ (strong). For immunofluorescence staining, YAP1 was co-stained with cytokeratin 5 (CK5) or 14 (CK14) (BioLegend, San Diego, CA) and imaged with a Nikon fluorescence microscope (Melville, NY).

### Gene silencing and luciferase assay

PCa cells were co-transfected with β-Catenin-responsive TOPflash plasmid and control or siRNAs against YAP1 (Santa Cruz Biotechnology) by using X-tremeGENE Transfection Reagent (Sigma Aldrich, St. Louis, MO). Luciferase activity was measured by using Promega Luciferase Assay kit (Promega Biotechnology, Maddison, WI).

### Western blotting

Equal amounts of protein were subjected to SDS-PAGE and then transferred to a PDVF membrane (Bio-Rad, Hercules, CA). Subsequently, membranes were incubated with primary antibodies and HRP-conjugated secondary antibody (Cell Signaling, Beverly, MA). Proteins were revealed by using ProSignal® Dura ECL Reagent (Genesee Scientific, San Diego, CA) and visualized in a Chemidoc™ Touch Imaging System (Bio-Rad).

### Statistical analyses

Differential gene expression was evaluated using Chi-Square and two-sided Student’s *t*-test. *p*-value <0.05 was considered statistically significant.

## RESULTS

### YAP1 expression in normal prostate

We used immunohistochemistry to assess the expression of YAP1 in benign prostatic tissues. YAP1 expression was detected in the nuclei of the prostatic p63-positive basal epithelial cells, as well as stromal cells, but it was absent in the p63-negative luminal glandular epithelial cells (Figs. 1A & 1B). Additionally, dual immunofluorescence staining of YAP1 and CK5 or CK14 that highlights prostatic basal cells was performed to corroborate the YAP1 co-localization pattern in benign basal prostatic epithelium. In concordance with previous data, YAP1 immunostaining was mainly present in the nuclei of the cells that were positive for CK5 and CK14 (Figs. 1C to 1H). These results indicate that the expression of YAP1 is localized in basal epithelial cells.

**Fig. 1.**
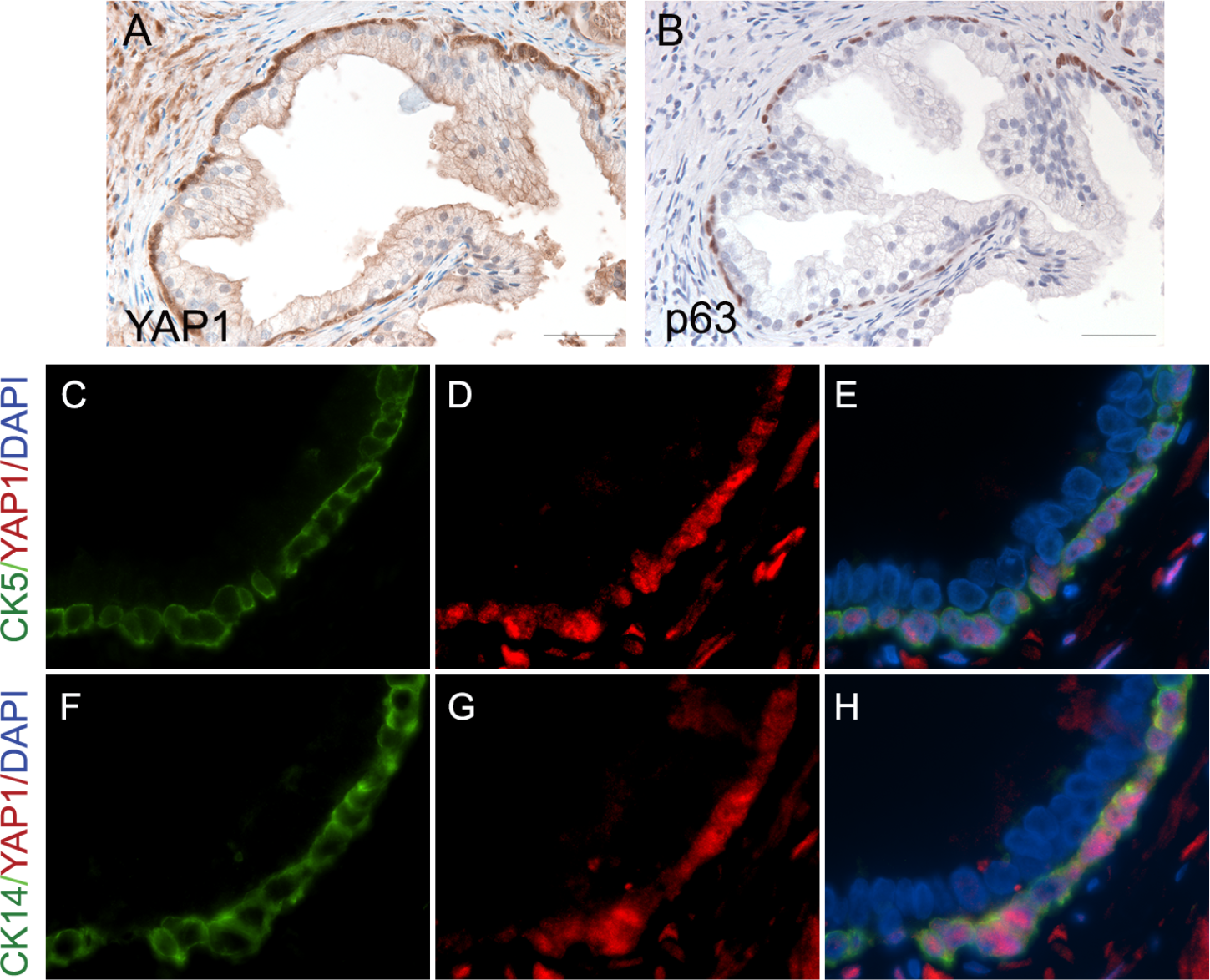
The expression of YAP1 in benign prostatic hyperplasia. (**A**) YAP1 displayed positive staining in basal cells (positive nuclear staining) and negative staining in the prostatic luminal epithelial cells. (**B**) Immunohistochemical staining of p63 highlighted basal cells in the periphery of the prostatic gland. (**C to H**) Dual immunofluorescence staining of YAP1 and cytokeratin 5 (CK5) or cytokeratin 14 (CK14) confirmed the presence of YAP1 expression in prostatic basal epithelial cells.

### Expression of YAP1 increased in PIN but lost in mouse prostate NEPC

We used TRAMP and LADY, the two widely used PCa mouse models,^15,16^ to evaluate YAP1 expression during PCa progression. TRAMP mice develop PIN and a subset of tumors progress into NEPC.^17^ We found that YAP1 expression was absent in the luminal epithelial cells in wild-type prostates of TRAMP mice (Fig. 2A, n=3) but present in PIN (Fig. 2B, n = 3). Rare clusters of PIN cells stained positive for FOXA2, a marker of NEPC (Fig. 2C). In the TRAMP tumors that developed NEPC (n=5), YAP1 was expressed in PIN lesions within the NEPC tumors (Fig. 2D). However, YAP1 was undetectable in the NEPC cells (Fig. 2D) which were highlighted by NEPC marker chromogranin A (CHGA) (Fig. 2E). The NEPC marker FOXA2 showed diffuse immunoreactivity in NEPC cells in contrast to its rare-to-negative stain in PIN tissues (Figs. 2C & 2F).

**Fig. 2.**
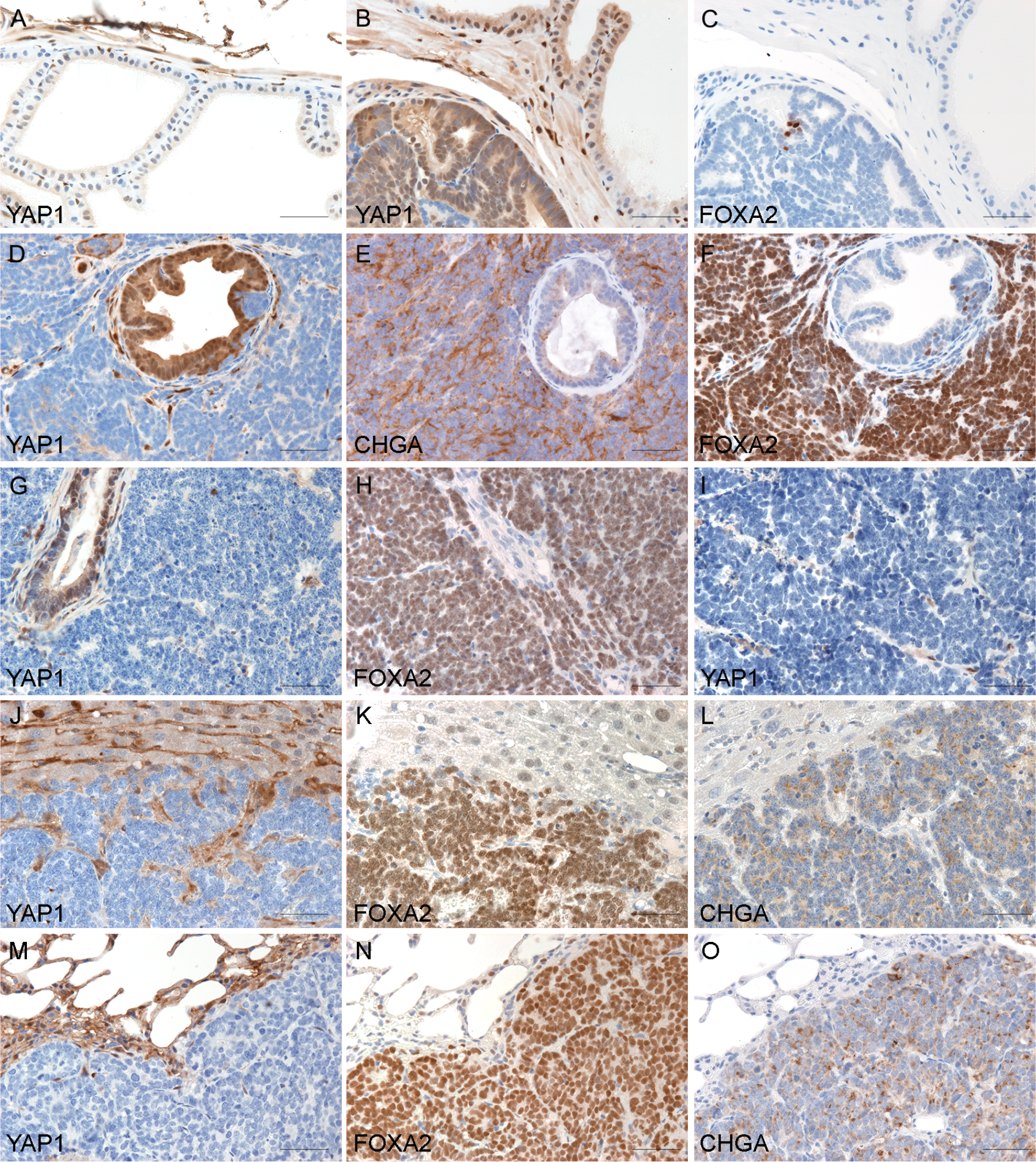
Microscopic examination of the prostate of TRAMP and LADY mice revealed that the expression of YAP1 increased in PIN but decreased in NEPC. (**A**) Wild-type prostate demonstrating positive staining of YAP1 in the occasional basal epithelia but negative staining in luminal epithelia. (**B&C**) Serial sections of a PIN tumor of TRAMP demonstrating diffuse positive staining of YAP1 and the rare FOXA2-positive cells in PIN. (**D to F**) Serial sections of a NEPC tumor of TRAMP demonstrating diffuse positive staining of YAP1 in PIN components but negative in NEPC cells. NEPC markers chromogranin A (CHGA) and FOXA2 were stained positive in NEPC cells but negative in PIN. (**G & H**) Serial sections of a 12T-10 NEPC tumor demonstrating negativity of YAP1 in NEPC cells in contrast to the positive staining in focal PIN. Positive staining of FOXA2 was seen in the NEPC cells. (**I**) NE10 tumor showed negative YAP1 staining. (**J** to **O**) The expression of YAP1 in NEPC metastases. Sections of the liver (J to L) and lung (M to O) with metastasis of NEPC demonstrated negative YAP1 staining in metastatic NEPC. In contrast, FOXA2 and CHGA were positive in NEPC metastases.

We also examined the expression of YAP1 in LADY mice including 12T-10, a NEPC model,^16^ and NE10, a xenograft line derived from 12T-10 NEPC.^18^ In 12T-10 LADY tumors, YAP1 expression was present in focal PIN lesions but was absent in the FOXA2-positive NEPC cells (Figs. 2G & 2H). YAP1 expression was lost in NE10 tumors (Fig. 2I). Additionally, YAP1 immunostaining was absent in NEPC metastases to the liver and lung, which expressed NEPC markers FOXA2 and CHGA (Figs. 2J to 2O). These results indicate that mouse-derived NEPC tumors exhibit a marked reduction in YAP1 expression compared to the non-NE tumors.

### The expression of YAP1 increased in human high-grade prostate adenocarcinomas but decreased in NEPC

YAP1 expression levels were also evaluated in human prostatic tissues including benign prostatic hyperplasia, low grade adenocarcinomas (AdPCa), high grade AdPCa, and NEPC. The results were summarized in suppl. Tables 1 & 2. YAP1 expression was detected in fibromuscular stromal cells in most cases. However, YAP1 expression in PCa was altered during progression of the disease (Figs. 3A to 3I). YAP1 was expressed in basal epithelial cells as well as fibromuscular stromal cells in benign prostatic tissues but absent from the luminal epithelial cells except of one case of benign prostatic glandular tissues (Fig. 3A, n=22). YAP1 expression was also absent in human high-grade PIN (2 of 2 cases), which was in contrast to the positive expression in mouse prostatic PIN. Of the 4 low-grade AdPCas with GS 6, 3 cases demonstrated absence of YAP1 expression in the acini of AdPCa (Fig. 3B) and only one tumor showed weak (1+) positive nuclear staining of YAP1. In AdPCa with GS 7, a subset of tumors (10 of 25 cases, 40%) exhibited positive YAP1 nuclear localization. Approximately 1-25% of cancer cells demonstrated various degrees of intensity (from 1+ weak to 3+ strong) of YAP1 staining. Of the 5 AdPCa with GS 8, 4 cases (80%) were positive, with one tumor showing strong YAP1 immunostain of 3+ in 50-75% of cancer cells and 2+ in the remaining cells. In AdPCa with GS 9, 10 of 14 (71%) tumors showed ~ 50-75% of cells with moderate (2+) to strong (3+) YAP1 immunostaining while 4 of 7 (57%) AdPCas with GS 10 exhibited as many as 3+ (50-75%) of cancer cells with 2+ (moderate) to 3+ (strong) YAP1 expression. Overall, the expression of YAP1 was associated with high-grade AdPCa (Gleason score≥8) (Chi-Square Test, *p* <0.05).

**Fig. 3.**
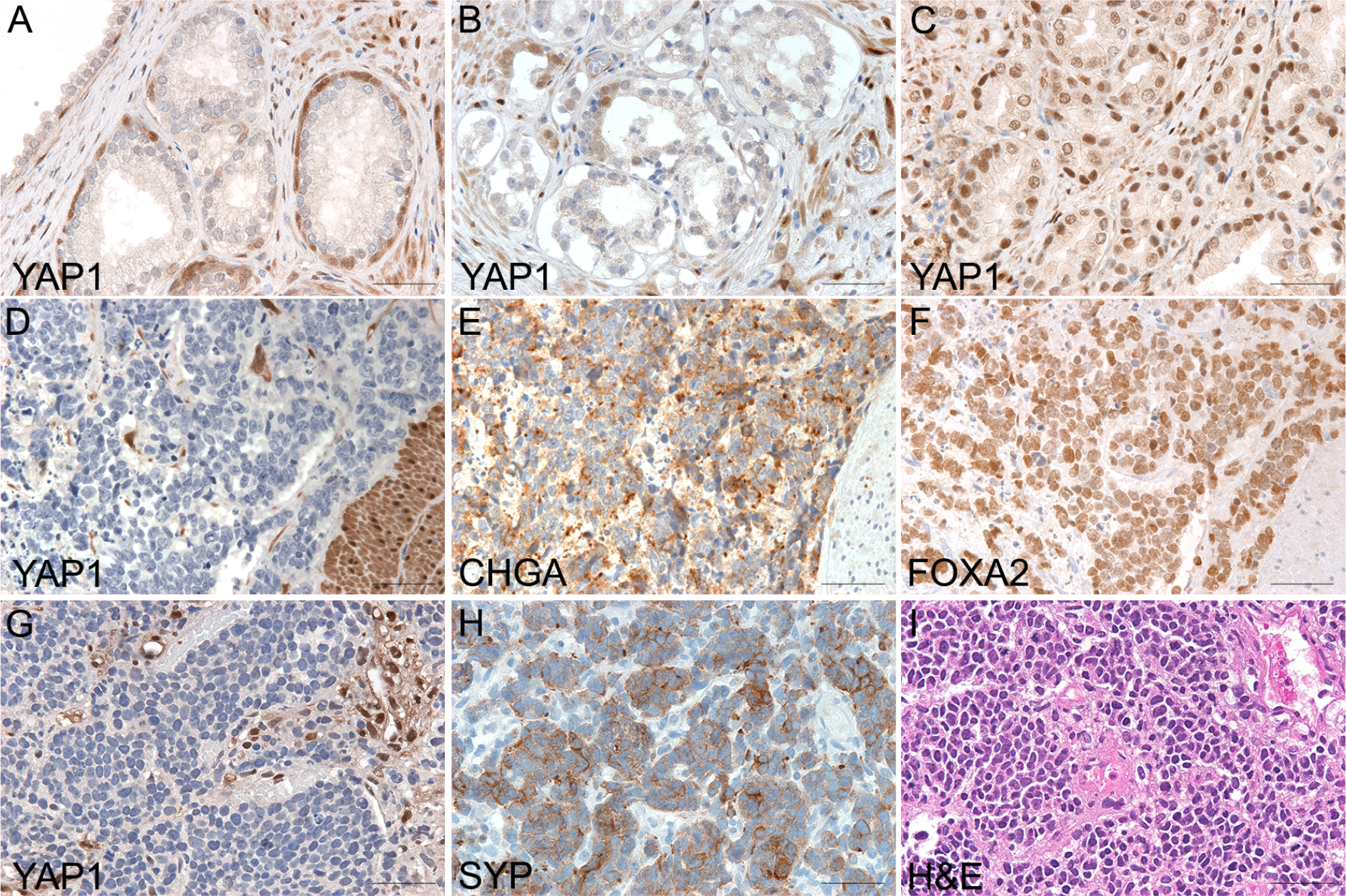
The expression of YAP1 in human prostatic tissues. (**A**) Benign prostatic tissue demonstrated positive YAP1 staining in the stroma cells and basal cells. (**B**) Low-grade prostate adenocarcinoma. YAP1 expression was present in stromal cells and occasional basal cells but was absent in adenocarcinoma cells. (**C**) High-grade prostate adenocarcinoma. YAP1 was expressed in prostate adenocarcinomas with a Gleason score of 8 or higher. (**D to F**) Serial sections of a NEPC with focal adenocarcinoma. YAP1 protein was expressed in high-grade adenocarcinoma components, but the expression was lost in NEPC cells. CHGA and FOXA2 were diffusely expressed in NEPC but not in adenocarcinoma cells. (**G to I**) Serial sections of a small cell carcinoma demonstrating negative YAP1 and positive synaptophysin (SYP) staining in NEPC cells.

**Table 1.** YAP1 expression in prostate adenocarcinoma. Note: ALL but one benign prostate specimens (n=22) displayed negative IHC staining of YAP1 in luminal epithelial cells.

|  | Tumors with positive YAP1 staining in cancer cells |
| --- | --- |
| GS 6 | 1/4 |
| GS 7 | 10/25 |
| GS 8 | 4/5 |
| GS 9 | 10/14 |
| GS 10 | 4/7 |

However, YAP1 expression was absent in 6/12 NEPC (Figs. 3D & 3G, suppl. Table 2, n=12), including one AdPCa with Paneth-like NE differentiation, one large cell NEPC, one unspecified NEPC and three small cell NEPC. In the NEPC with focal adenocarcinoma (Fig. 3D), YAP1 expression was present in the adenocarcinoma cells but absent in NEPC components. Chromogranin A (CHGA) (Fig. 3E), FOXA2 (Fig. 3F), and Synaptophysin (SYP) (Fig. 3H) were used as markers for NEPC. These results demonstrate that YAP expression is lost in NEPC.

**Table 2.** YAP expression in neuroendocrine prostate cancer

|  |  | Percentage of YAP1<sup>+</sup> cells | Intensity of staining |
| --- | --- | --- | --- |
| PCIM 41 | Paneth cell | 0 | 0 |
| Prostate 7 | NEPC | 0 | 0 |
| PCIM 1 | large cell | 0 | 0 |
| Prostate 4 | small cell | 0 | 0 |
| Prostate 12 | small cell | 0 | 0 |
| Prostate 14 | small cell | 0 | 0 |
| Prostate 2 | small cell | <25% | 3+ |
| Prostate 5 | small cell | <25% | 1+/2+ |
| Prostate 8 | small cell | <25% | 1+/2+ |
| Prostate 9 | small cell | <25% | 1+/2+ |
| Prostate 10 | small cell | <25% | 1+/2+ |
| Prostate 11 | small cell | <25% | 1+/2+ |

To further validate the reduction of YAP1 expression in NEPC, we analyzed mRNA levels of YAP1 in an RNA-seq dataset of 46 LuCaP, PCa patient-derived xenograft models (PDXs). The expression levels of YAP1, TAZ, AR, and NEPC markers including FOXA2, SOX2, SYP and CHGA were displayed as a heatmap. As shown in Fig. 4A, YAP1 expression was lost in all 8 NEPC LuCaP PDXs as well as the amphicrine LuCaP 77CR line that is AR^positive^ NE^positive^. However, loss of YAP1 expression only occurred in 1 of 35 AR^positive^ AdPCa. YAP1 expression was detectable in the double negative LuCaP 173.2 as well as the AR^low^/NE^negative^ LuCaP 176 PDX. In contrast to the reduced YAP1 expression in NEPC, TAZ expression in these LuCaP PDXs did not show any consistent patterns (Fig. 4). Statistical analysis revealed that the loss of YAP1 expression correlated with NE phenotype (Chi-Square test, *p*<0.001). Taken together, these data further support the lack of expression of YAP1 in NEPC.

**Fig. 4.**
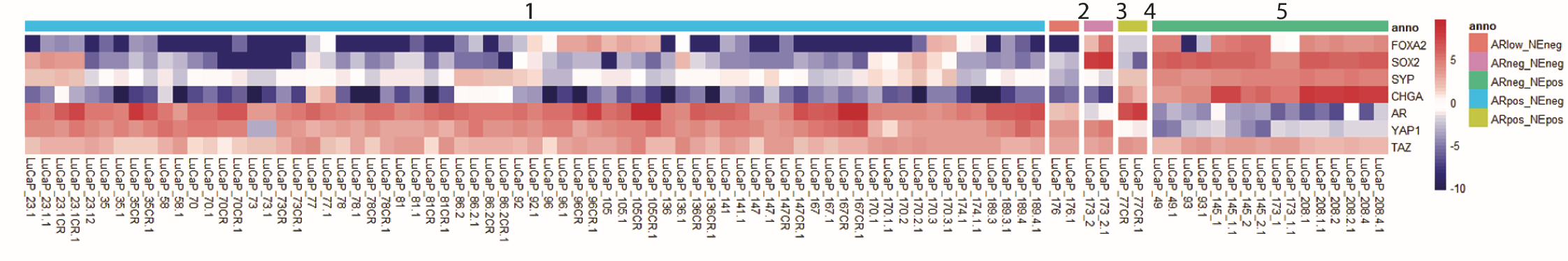
The expression of YAP1 in LuCaP PDXs. The expression levels of YAP1, TAZ, AR, and NEPC markers (FOXA2, SOX2, CHGA, and SYP) were extracted from RNAseq data derived from 46 LuCaP PDXs including (from left to right) AR^+^/NE^−^ (group 1, n=35), AR^low^/NE^−^ (group 2, n=1), AR^−^/NE^−^ (group 3, n=1), AR^+^/NE^+^ (group 4, n=1), and AR^−^/NE^+^ tumors (group 5, n=8). YAP1 expression was lost in all the NE positive LuCaP PDXs. Whereas, YAP1 was expressed in all but one prostate adenocarcinoma cases.

### YAP1 loss in AdPCa cells induces Wnt/β-Catenin activity

Hippo/YAP1 pathway has a close crosstalk with Wnt/β-Catenin signaling,^7,8^ and previous studies have linked the activation of Wnt/β-Catenin signaling with NE differentiation of PCa.^5,12^ Therefore, we assessed the effects of YAP1 knockdown on the Wnt/β-Catenin activity. The activity of Wnt/β-Catenin increased in PCa PC3 cells (Fig. 5) as well as in 22Rv1 cells (data not shown) after YAP1 knockdown. However, YAP1 knockdown did not cause significant changes in the expression of NEPC markers (data not shown). Taken together, these data indicate that the downregulation of YAP1 activates Wnt/β-Catenin signaling in AdPCa cells.

**Fig. 5.**
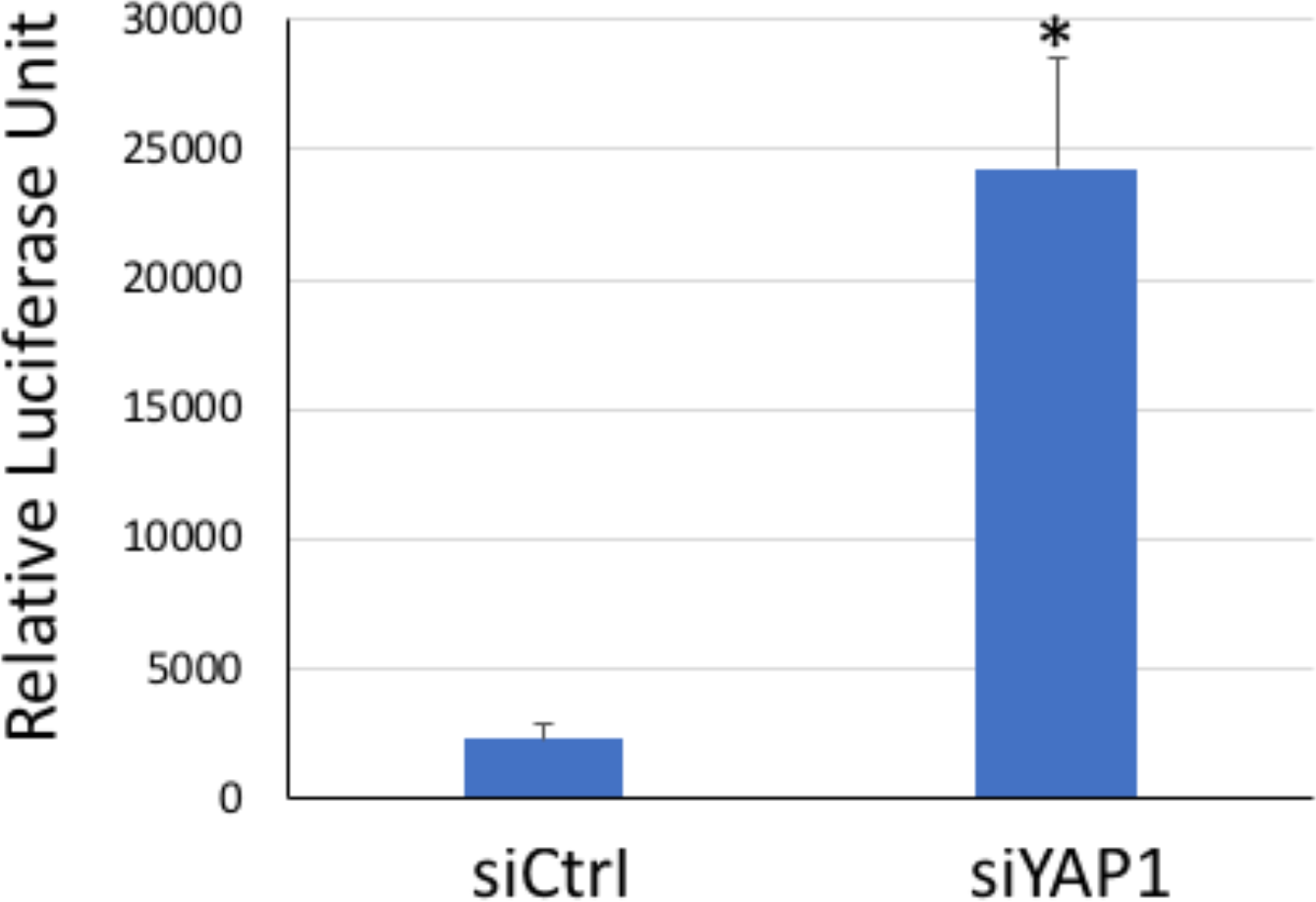
YAP1 knockdown induces Wnt/β-Catenin activity. Luciferase assay was conducted using PCa cells that were co-transfected with TOPFlash plasmid and siYAP1. YAP1 knockdown induced Wnt/β-Catenin activity in PCa cells (*t*-test, *p*<0.05).

## DISCUSSION

The development of NEPC is a clinically important issue. We need to elucidate the pathogenesis of NEPC and develop an effective therapy for this aggressive disease.^3^ In this study, we reported that the Hippo pathway coactivator YAP1 was reduced/lost in NEPC, and a reduction of YAP1 provided a mechanism to activate Wnt/β-Catenin signaling in PCa cells. Given the implication of Wnt/β-Catenin signaling in NEPC progression,^5,12^ these results suggest a role of YAP1 in the pathogenesis of NEPC.

In cancer, YAP1 usually functions as an oncogene, promoting cell proliferation and invasion.^6^ However, YAP1 also exerts tumor suppressive role in certain cellular contexts. For example, YAP1 overexpression reduces small cell lung carcinoma (SCLC) growth.^21^ Inversely, its loss stimulates breast cancer cell proliferation and mobility, protecting them from anoikis.^22^ These data suggest that YAP1 exerts a dual-role during tumorigenesis. Therefore, it would be interesting to monitor the changes in its expression during PCa progression. In this study, we demonstrated that the expression of YAP1 was limited to the basal cells in benign prostatic glandular epithelia and was absent in the luminal glandular cells, supporting the previous proposal that YAP1 may serve as a biomarker for prostatic basal epithelial cells.^23^ YAP1 expression was absent in the cancer cells of early stage AdPCa where there were no basal cells in the glandular epithelium. Its expression was increased; however, in advanced stages of AdPCa. The increased levels of YAP1 expression was associated with high-grade AdPCa (Fig. 3) and advanced tumor-node-metastasis (TNM) stages.^24,25^ Additionally, the inhibition of this protein in high-grade PCa cells not only impaired cellular proliferation, but also enhanced cellular sensitivity to ADT, ^11,26^ supporting that inhibition of YAP1 could control AdPCa progression.

However, the expression of YAP1 was decreased/lost in cell line, human, and mouse NEPC tumor samples (Figs. 2 to 4). These findings were consistent with that demonstrated in SCLC cells,^10,27^ where the YAP1 expression is reduced in SCLC and its downregulation is essential to NE differentiation.^10,27^ However, the molecular mechanism that relates the loss of YAP1 with the acquisition of NE phenotype has not been elucidated. Prior studies in lung cancer have demonstrated that YAP1 knockdown induces the expression of several neuroendocrine markers such as Rab3A to achieve NE differentiation.^10^ In this study, we showed that downregulation of YAP1 activated Wnt/β-Catenin signaling, and the activation of Wnt/β-Catenin signaling has been previously related to the induction of NE differentiation of PCa.^5,12^ Additionally, previous research has shown that YAP1/TAZ inhibit the activation of Wnt/β-Catenin signaling by recruiting β-TrCP to the β-Catenin destruction complex, an essential step for inducing β-Catenin’s degradation by the ubiquitin-proteasome system.^7^ Consequently, the downregulation of YAP1 could induce the nuclear translocation of β-Catenin resulting in the activation of Wnt/β-Catenin signaling.^8^ In consistent with these findings^28,29^, our studies demonstrated that Wnt/β-Catenin activity increased after YAP1 knockdown, supporting the idea that loss of YAP1 provides a mechanism to activate Wnt/β-Catenin signaling in AdPCa cells. However, transient YAP1 knockdown was not sufficient to induce the expression of NEPC markers or the NE-like cell morphology. More studies would follow to determine whether long-term loss of YAP1 could promote NE differentiation of AdPCa cells.

Although the levels of YAP1 decreased/lost in NEPC, the closely-related TAZ did not display consistent changes in prostatic tissues (Fig. 4). Despite the high homology of YAP1 and TAZ proteins, they have different structures, are involved in differential protein-protein interactions, and have different patterns of downstream regulations. In some cellular processes, they even exhibit antagonistic physiologic responses.^30,31^ Our results support that YAP1 and TAZ may have distinct roles in NEPC.

Taken together, our results highlighted the alteration of YAP1 expression during PCa progression. Although previous studies have linked YAP1’s overexpression with increased AdPCa malignancy, we showed here its downregulation activates the Wnt/β-Catenin signaling, which could lead to NE differentiation. However, more studies would be necessary to determine how the loss of YAP1 affects NE differentiation, as well as if YAP1 inhibitors could be safely used as therapeutic adjuvants for treating AdPCa.

## ACKNOWLEDGMENT

We thank Dr. Robert Matusik at Vanderbilt University for providing tissues of LADY mice and advice on this research. This research was supported by NIH R03 CA212567, R01 CA226285, U54 GM104940, DOD W81XWH-12-1-0212, and LSUHSC FWCC and Office of Research funding to Yu, X.

## CONFLICT OF INTEREST

We declare no conflicts of interest.

